# Blimp-1 mediates tracheal lumen maturation in *Drosophila melanogaster*

**DOI:** 10.1101/273151

**Authors:** Arzu Öztürk-Çolak, Camille Stephan-Otto Attolini, Jordi Casanova, Sofia J. Araújo

## Abstract

The specification of tissue identity during embryonic development requires precise spatiotemporal coordination of gene expression. Many transcription factors required for the development of organs have been identified and their expression patterns are known; however, the mechanisms through which they coordinate gene expression in time remain poorly understood. Here we show that hormone-induced transcription factor Blimp-1 participates in the temporal coordination of tubulogenesis in *Drosophila melanogaster* by regulating the expression of many genes involved in tube maturation. In particular, we demonstrate that Blimp-1 regulates the expression of genes involved in chitin deposition and F-actin organization. We show that Blimp-1 is involved in the temporal control of lumen maturation by regulating the beginning of chitin deposition. We also report that Blimp-1 represses a variety of genes involved in tracheal maturation. Finally, we reveal that the kinase Btk29A serves as a link between Blimp-1 transcriptional repression and apical extra-cellular matrix organization.

## Introduction

Specialized cellular functions and cell lineage fates are usually regulated by only a few key instructive transcription factors required to activate or repress specific patterns of gene expression. During the development of multicellular organisms, these events must be precisely timed. As a result, organogenesis requires an accurate spatio-temporal regulation of gene expression over extended periods. While many transcription factors required for the development of organs have been identified and their expression pinpointed spatially, the mechanisms through which they coordinate downstream gene expression in time remain poorly understood. Here we addressed part of this question during the development of the *Drosophila melanogaster* tracheal system, a model used to study epithelial organ development. The tracheal system of *D. melanogaster* is formed by a network of epithelial tubes that requires tight temporal regulation of gene expression. Tracheal tube maturation involves the timely and spatially regulated deposition of a chitinous apical Extracellular Matrix (aECM), a process that is governed by downstream effectors of the mid-embryonic ecdysone hormone pulse (Chavoshi *et al.* 2010). One of these ecdysone response genes is the *D. melanogaster B-Lymphocyte Inducing Maturation Protein-1 (Blimp-1)* (Ng *et al.* 2006; Chavoshi *et al*. 2010). *Blimp-1* is the homolog of human *Prdm1 (Positive regulatory domain containing 1)* (Huang 1994). Blimp- 1/PRDM1 is a zinc finger transcriptional repressor that belongs to the Prdm gene family and it was originally identified as a silencer of *β-interferon* gene expression (Keller and Maniatis 1991). Prdm family members contain a conserved N-terminal domain, known as a positive regulatory domain (PR domain). This domain has been associated with the SET methyltransferase domain, which is important for the regulation of chromatin-mediated gene expression (Hohenauer and Moore 2012). In addition, Prdm family proteins contain multiple zinc fingers that mediate sequence-specific DNA binding and protein-protein interactions (Turner *et al.* 1994). Prdm family members modulate key cellular processes, including cell fate, and the aberrant function of some members may lead to malignant transformation (Fog *et al.* 2012). During embryonic development, Blimp-1 controls a plethora of cell-fate decisions in many organisms (Bikoff *et al.* 2009; Hohenauer and Moore 2012). In *D. melanogaster, Blimp- 1* serves as an ecdysone-inducible gene that regulates *ftz-f1* in pupal stages (Agawa *et al.* 2007). By acting as a transcriptional repressor, *Blimp-1* prevents the premature expression of *ftz-f1*, thereby influencing the temporal regulation of events that are crucial for insect development. The expression level and stability of Blimp-1 is critical for the precise timing of pupariation (Akagi *et al.* 2016).

*Blimp-1* exerts a function in tracheal system morphogenesis during embryonic development (Ng *et al.* 2006; ÖztÜrk-Çolak *et al.* 2016). However, the question remains as to how this transcription factor regulates tube maturation events downstream of the hormone ecdysone. Here we studied the role of Blimp-1 in the transcriptional regulation of the regulation of tracheal tube maturation in *D. melanogaster.* We found that Blimp-1 is a transcriptional repressor of many genes involved in tracheal development and that its levels are critical for the precise timing of luminal maturation and the final stages of tubulogenesis in the embryo. Our results indicate that Blimp-1, working downstream of ecdysone, acts as a link of hormone action during tube maturation in organogenesis.

## Materials and Methods

### *D. melanogaster* strains and genetics

All *D. melanogaster* strains were raised at 25°C under standard conditions. Mutant chromosomes were balanced over LacZ or GFP-labelled balancer chromosomes. Overexpression and rescue experiments were carried out either with *btl-GAL4* (kindly provided by M. Affolter) or *AbdB-GAL4* (kindly provided by E. Sánchez-Herrero) drivers at 25°C or 29°C. *y^1^w^118^* (used as wild-type), *Blimp-1^KG09531^*, and *UAS-srcGFP* are described in FlyBase; UAS- Blimp-1 (ÖztÜrk-Çolak *et al.* 2016); Btk29A^k00206^ and UAS-Btk29A (kindly provided by M. Strigini).

### Embryo staging and synchronization

Embryos were staged following (Campos-Ortega and Hartenstein 1985). To study temporal chitin deposition, cages were set at 25°C for 2 h, and embryos were then allowed to develop for 18 h, 20 h, 22 h or 24 h at 18°C in order to obtain early and late stages 12, and early and late 13, respectively. *Blimp-1* mutant embryos were compared to *Blimp-1* heterozygotes after staining with CBP and 2A12. Heterozygote embryos were differentiated from homozygous mutant embryos by the presence or absence of a β-Gal-expressing balancer.

### Synthesis of *pri/tal* RNA probes for in-situ hybridization

The *pri/tal* RNA probes were synthesized using a PCR-based technique. The *pri* gene region (524bp, covering all coding Open Reading Frames (ORFs) of the gene) was defined, and the forward (5’TAATACGACTCACTATAGGTTTTTGGTCAATACACGGCA3’) and reverse (5’AATTAACCCTCACTAAAGGAGTTTGTGGATAAGGCACGG3’) primers were designed accordingly so that the PCR product carried the two RNA promoters T3 and T7. The gene region of interest, flanked by the T3 and T7 sequences, was amplified from previously isolated genomic DNA via PCR under standard PCR conditions. The newly synthesized RNA was then purified by precipitation, re-suspended in hybridization buffer, and stored at -20°C.

### Immunohistochemistry, image acquisition and processing

Standard protocols for immunostaining were applied. The following antibodies were used: rat anti-DE-cad (DCAD2, DSHB); rabbit anti-GFP (Molecular Probes); anti-Gasp mAb2A12 (DSHB); guinea pig anti-Blimp-1(S. Roy); anti-expansion and anti-rebuf (from M. Llimargas); anti-knk (from A. Uv) anti Btk29A (from M. Strigini); anti aPKC (Santa Cruz Biotechnology); and chicken anti-β-gal (Cappel). Biotinylated or Cy3-, Cy2- and Cy5-conjugated secondary antibodies (Jackson ImmunoResearch) were used at 1:300. Chitin was visualised with Fluostain (Sigma) at 1 μg/ml or CBP (Chitin Binding Probe, our own, made according to NEB protocols). Confocal images of fixed embryos were obtained either with a Leica TCS-SPE, a Leica TCS- SP2, or a Leica TCS-SP5 system. Images were processed using Fiji and assembled using Photoshop. 3D cell shape reconstructions were done using Imaris software.

### Fluorescent in-situ hybridization

Freshly fixed embryos were washed and kept at 56°C in Hybridization Buffer for 3 h for pre-hybridization. In the last 10 min of pre-hybridization, probes (1:100 in Hybridization Buffer) were prepared for hybridization. The probes were hybridized with the embryos at 56°C overnight. The next day the embryos were washed and incubated in POD-conjugated anti-Dig (in PBT) for 1 h. The fluorescent signal was developed by the addition of Cy3 Amplification Reagent (1:100) diluted in TSA Amplification Diluent and incubation at room temperature in the dark for 10 min. Finally, the embryos were either mounted in Fluoromount medium or subjected to antibody staining.

### In silico analysis of Blimp-1 binding sites

The Blimp-1 position weight matrix was taken from the reported binding sequences in (Kuo and Calame 2004). We extracted sequences 2000 bp up- and down-stream from all annotated isoforms in the *D. melanogaster* genome using the biomaRt database (Smedley *et al.* 2015). All computations were performed within the R statistical framework (http://www.R-project.org). The Matscan software (Blanco *et al.* 2006) was used to find putative binding sites for the Blimp-1 position weight matrix in the aforementioned regions. For each binding site, we computed the mean conservation score of the corresponding positions following the *D. melanogaster* related species’ conservation track of the USCS browser (Fujita *et al.* 2011). For genes with multiple transcription start sites (TSSs), non-redundant binding candidates for all TSSs were reported. Each gene was assigned with the maximum Matscan score of the corresponding binding sites after filtering by conservation score.

## Results

### Blimp-1 modulates tracheal tube size and apical ECM formation

*Blimp-1*, an ecdysone response gene (Beckstead *et al.* 2005; Chavoshi *et al.* 2010) (Supplementary Fig. S1), encodes the *D. melanogaster* homolog of the transcriptional factor *B-lymphocyte-inducing maturation protein* gene, whose mutants have been reported to have misshapen trachea with severe defects in taenidia (Ng *et al.* 2006; ÖztÜrk-Çolak *et al.* 2016). Detailed expression analysis of Blimp-1 protein in tracheal cells showed that expression is not detectable until embryonic stage 12 and is then observed until stage 15 (Fig.1 A-D). At this stage, Blimp-1 protein levels started decreasing, first in dorsal trunk (DT) cells and by stage 16 the protein was no longer detectable in any tracheal cell. In addition to its tracheal expression, Blimp-1 was also detected in the epidermal, midgut and hindgut cells throughout these stages (Fig.1 A-D). In *Blimp-1* mutant stage 16 embryos, we detected lower levels of chitin, inflated tracheal tubes and disorganized taenidial ridges (ÖztÜrk-Çolak *et al.* 2016) (Fig.1 E-H).

**Figure 1.**
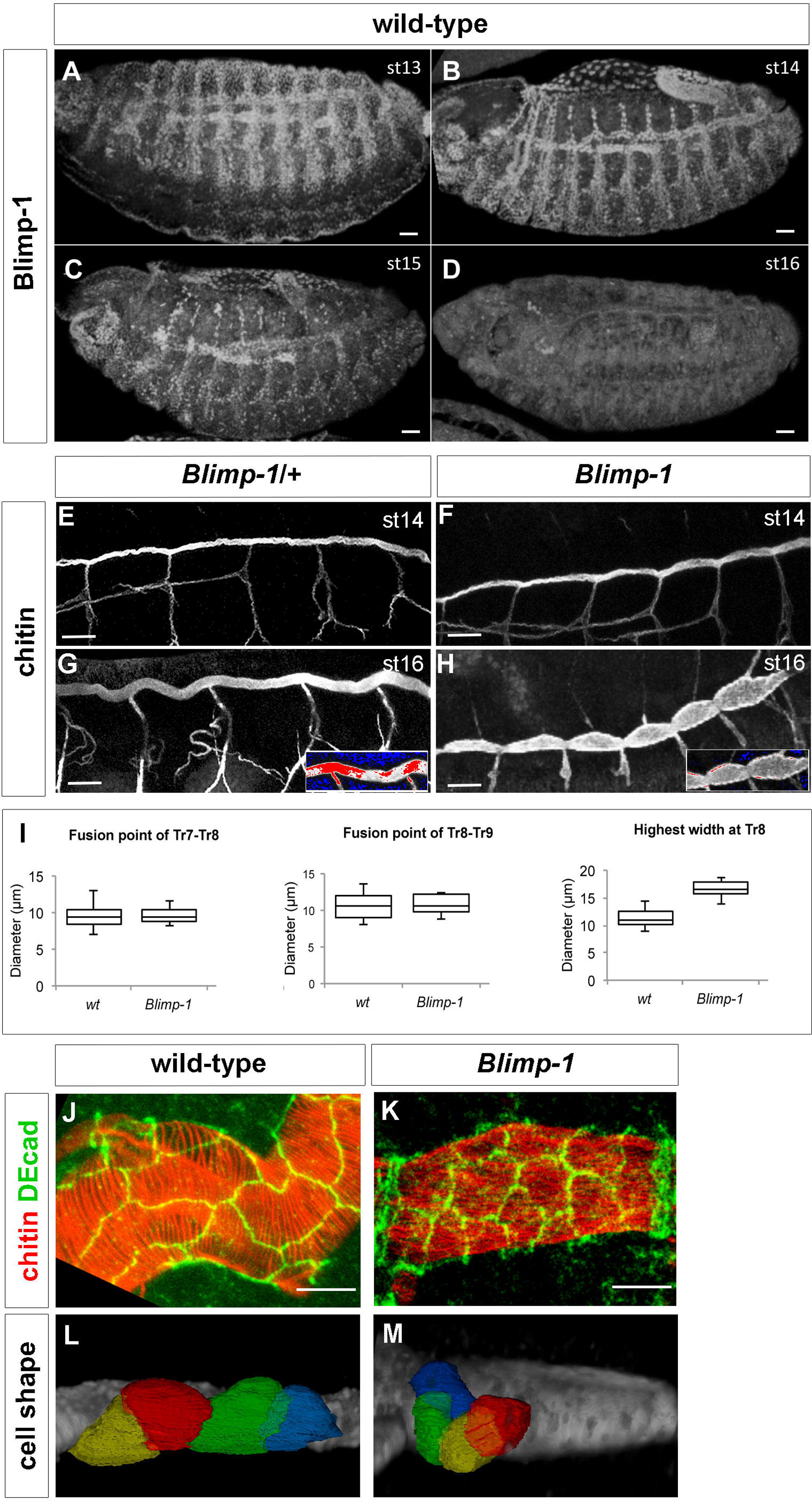
Blimp-1 is required for tracheal tube maturation. (A-D) *Wild-type* embryos stained for Blimp-1 protein with an anti-Blimp-1 antibody and showing Blimp-1 localization in tracheal cells from stage 13 to stage 15. At stage 15, Blimp-1 starts to be downregulated from the dorsal trunk (DT) and protein is not detected in tracheal cells at stage 16. Scale bars are 20 μm. (E-H) At stage 14, *Blimp-1* tracheal DT is not distinguishable from the *wild-type* DT. Differences are observed at later stages of development, when *Blimp-1* embryos display DTs with tube expansions between DT fusion points (H). (G and H) insets show detail of DT under a HiLo LUT, where highest intensity levels are red and lowest are blue, so differences in chitin amounts can be observed. Scale bars are 10 μm. (I) Quantification of tube width at fusion points and at between fusion points. (J-M) *Blimp-1* DT tracheal cells have smaller apical domains than *wild-type* DT tracheal cells. Scale bars are 5 μm.

The tube shape of *Blimp-1* mutant embryos was altered, DTs showing a smaller diameter at the fusion points than in the rest of the tubes (Fig.1 H). To better study whether this phenotype was due to constrictions at fusion points and/or dilations along the entire length of the tubes, we measured the tube diameter at fusion points of tracheal metameres Tr7-Tr8 and Tr8-Tr9 and the largest tube diameter at the Tr8 metamere between fusion points. *Blimp-1* mutant embryos had a significantly (p-value: 4.4E-07, by Student’s T-test, n=10) larger tube diameter between fusion points than *wild-type (wt)* DTs (n=15), while at the fusion points the tube diameter was similar to the *wt* (Fig.1 I I). These results suggest that *Blimp-1* is required to maintain the luminal structures from over-expanding in between fusion points.

Since tube expansion is related to the apical cell surface, we next examined cell shape in the trachea of *Blimp-1* mutant embryos. We labelled the apical cell junctions in *Blimp-1* mutants and found that the apical cell shape differed to that of *wt* cells. In *Blimp-1* mutants, the longest cell axis appeared to be perpendicular to the tube axis, while in the *wt* it was parallel (Fig.1 J-M). Most of the *Blimp-1* mutant cells had a similarly reduced apical cell surface area, thereby suggesting a uniform organization throughout the tube (except for the fusion cells, which already had a distinct apical cell shape that did not seem to be affected by the loss of function *of Blimp-1)* (Fig. 1 K). To have a better idea of cell organization in *Blimp-1* mutant trachea, we traced tracheal cell surfaces and compared them to those of the *wt.* We observed altered cell shapes in the former. In this regard, the mutant trachea showed cells that were less elongated and more “square-shaped” than *wt* ones (Fig.1 L, M).

Taken together, these observations reveal that Blimp-1 affects various stages of tube maturation, from chitin deposition to tube expansion, as well cellular morphology.

### Blimp-1 modulates the timing of chitin deposition

In *wt* embryos, a matrix composed of chitin and proteins such as Gasp, accumulates in the tracheal lumen. This matrix plays a key role in the regulation of tube length and diameter expansion (Moussian *et al.* 2005; Tonning *et al.* 2005; Moussian *et al.* 2015). Due to the striking tube size phenotype *of Blimp-1* mutants, and because mutations in genes involved in chitin biogenesis and assembly result in irregular diametric expansion leading to locally constricted and dilated tubes, we examined the deposition of these markers from stage 12 (Fig. 2). In the wt, early chitin deposition began at stage 13, when a chitinous filament started to be deposited inside the DT just prior to tube expansion (Moussian *et al.* 2015). In parallel, Gasp was detected from stage 13, but at this stage it was mainly cytoplasmatic. None of these markers were detected at earlier stages. At stage 13, chitin began to be deposited in the lumen of all branches, starting from the DT (Moussian *et al.* 2015), and a mature pattern was achieved at stage 17, when taenidial ridges are fully formed (TiklovÁ *et al.* 2013; ÖztÜrk-Çolak *et al.* 2016). *Blimp-1* mutant embryos showed chitin deposition as early as stage 12 (Fig. 2 B, C). The pattern of chitin deposition in mutants at stage 13 was similar to that of the *wt* (Fig. 2 D, F). However, at later stages, the trachea of *Blimp-1* mutants showed lower levels of chitin than the *wt* (Fig. 2 G) (ÖztÜrk-Çolak *et al.* 2016). Gasp localization was lower in *Blimp-1* mutants throughout embryogenesis, especially in the DT during later stages (Fig. 2 K), a pattern resembling that of embryos mutant for genes involved mutants involved in chitin synthesis and organization (AraÚjo *et al.* 2005). We have previously shown that when we overexpressed *Blimp-1* in the posterior part of the embryo using Abd-BGAL4 to create tracheal DTs with distinct cellular compositions (FÖrster *et al.* 2009), we could detect lower levels of chitin in the posterior domain expressing higher levels of *Blimp-1* (öztürk-Qolak *et al.* 2016). To further analyse the influence of Blimp-1 in timely chitin deposition, we used the same experimental conditions and found that chitin was hardly detected in the posterior part of the trachea (Fig. 2 L) (Ozturk-Colak *et al.* 2016). However, further examination with higher laser power and gain showed that indeed both the chitin filament and taenidia were present throughout the trachea, although they displayed much lower levels in the posterior metameres, resembling chitin levels at earlier stages of development (Fig. 2 M). *Blimp-1* overexpression lead to a tubular structure that regarding chitin composition and organization was apparently younger than the same tube with normal levels of Blimp-1 protein. Thus, Blimp-1 seems to regulate the timing of the beginning of chitin deposition by repressing target proteins. When Blimp-1 levels are high, chitin deposition is delayed, whereas when they are low, chitin deposition begins earlier in the tracheal tubes.

**Figure 2.**
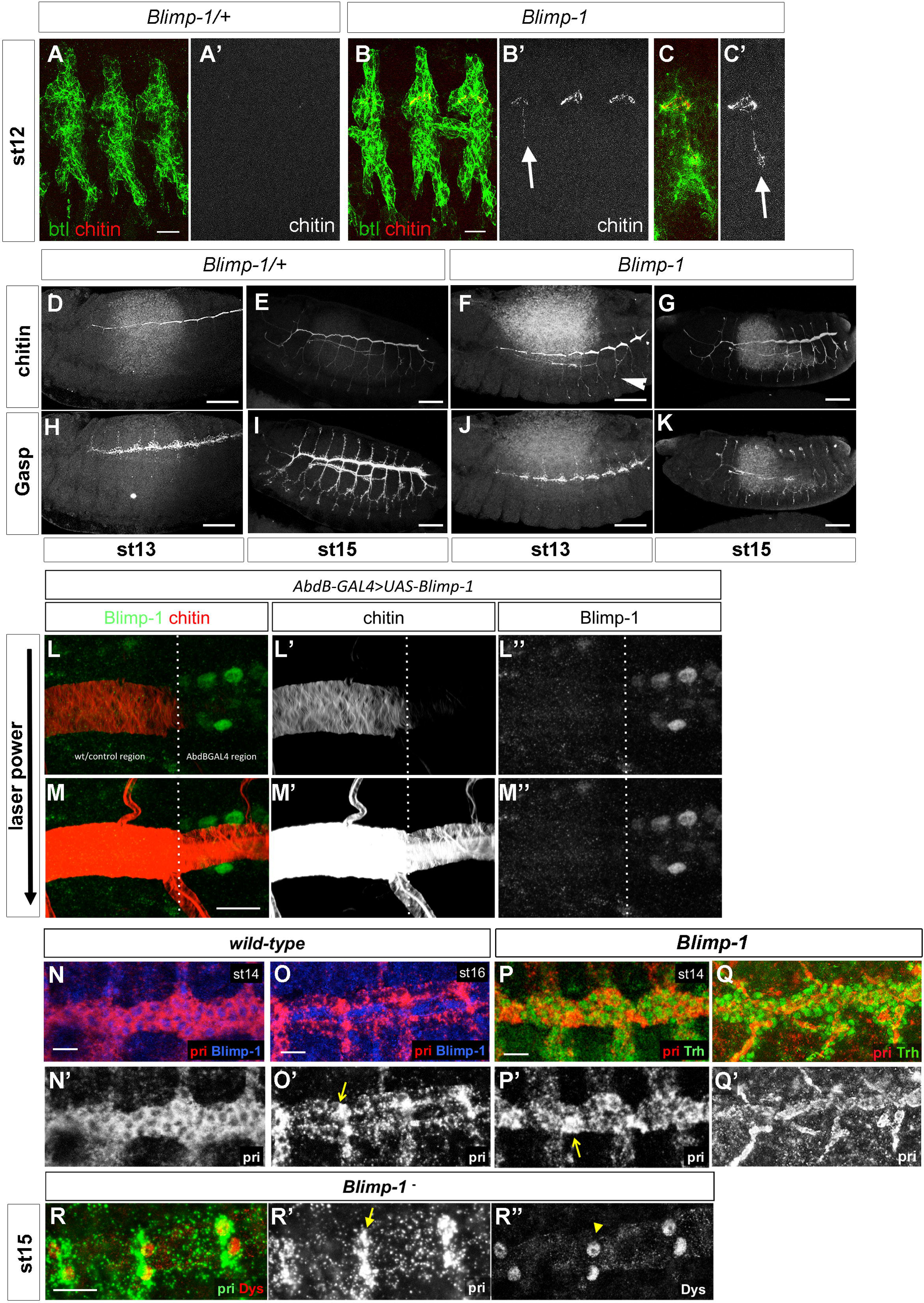
The timing of tube maturation is altered in *Blimp-1* mutants. (A-C) The deposition of chitin in *Blimp-1* embryos starts earlier than in the *wild-type* and can be detected in the primordia of the DT and in some of the transverse connectives (LTs, arrows) from stage 12 (B, C *vs.* A). Btl positive cells are detected by a btl::moeGFP construct, stained with GFP. Scale bars are 5 μm. (D-K) Comparison of chitin and Gasp deposition in heterozygous (D, E, H, I) vs. homozygous (F, G, J, K) *Blimp-1* embryos. Gasp deposition is defective in *Blimp-1* embryos at stage 15, regardless of earlier chitin deposition. Scale bars are 40 μm. (L, M) Detail of an early stage 17 DT at the border between cells expressing normal *Blimp-1* levels and higher Blimp-1 levels driven by AbdBGAL4. Overexpression of *Blimp-1* delays tube maturation, as reflected by lower chitin deposition. Scale bars are 10 μm. (N-P) Fluorescent *in-situ hybridization* of *pri* (red) co-stained with either anti-Blimp-1 (blue) to label Blimp-1-expressing cells, or anti-Trh (green) to label tracheal cells, in *wild-type* stage 14 (N) and stage 16 (O) and *Blimp-1* mutant stage 14 (P) embryos. In *wild-type* embryos, at stage 14 *pri* is uniformly expressed throughout the tube (N’). Later, at stage 16, *pri* expression starts to gradually disappear except at fusion points (O’, arrow). In the *Blimp-1* mutant embryo this differential expression of *pri* is already present at stage 14. Note the lower levels of *pri* expression throughout the tube at stage 14 of the *Blimp-1* mutant embryo (P), except at the fusion points (arrow). Scale bars are 10 μm. (Q) pri-expressing cells in *Blimp-1* loss of function correspond to the fusion cells. Fluorescent *in-situ hybridization* of pri (green) co-stained with anti-Dys (red) to label fusion cells in a *Blimp-1* mutant embryo at stage 15. The *pri* expression (arrow in Q’) co-localizes with *dys* expression (arrowhead in Q’’), indicating that the *pri-*expressing cells are fusion cells.

### The expression pattern *of tarsaless/polished* rice is altered in *Blimp-1* mutants

Mutants for *tarsal-less* (tal), also known as *polished rice* (pri), are affected in both F-actin and taenidia organization and we analysed them in parallel to *Blimp-1* mutants (ÖztÜrk-Çolak *et al.* 2016). Like *Blimp-1*, the tracheal expression of *tal/pri* in wild-type embryos is regulated by ecdysone (Chanut-Delalande *et al.* 2014)(Supplementary Fig. S1) and started in tracheal cells at stage 12 (Supplementary Fig. S2). At early stages this expression was restricted to DT cells but was later on observed uniformly in all tracheal cells until stage 15 (Supplementary Fig. S2). From this stage, *tal/pri* expression gradually decreased in all DT cells, except fusion cells, and at stage 16, coinciding with the time Blimp-1 disappears from tracheal cells, it became higher in fusion cells than in the rest of the DT (Supplementary Fig. S2 and Fig. 2 O). *tal/pri* expression in *Blimp-1* mutants at stage 14 was not as uniform as in the *wt*, as it was higher in some cells than in others (Fig. 2 P). To verify whether the cells with higher *tal/pri* expression were indeed fusion cells, we double labelled the embryos with the fusion cell marker *dysfusion (dys)* (Jiang and Crews 2003). High levels of both *tal/pri* and *dys* were detected in the same cells, thereby confirming that these were indeed fusion cells (Fig. 2 Q). These results indicate that *tal/pri* expression at stage 14 Blimp-1 embryos resembles expression of *tal/pri* in stage 16 *wt* embryos, suggesting that *Blimp-1* regulates the timing of the onset of *tal/pri* differential expression, from embryonic stage 16. We hypothesize that Blimp-1 might be involved in the differential regulation of *tal/pri* expression in tracheal cells. Blimp-1 could play a role in repressing *tal/pri* expression at high levels in fusion cells and, from stage 16, when Blimp-1 is no longer present in tracheal cells, this repression would no longer be exerted leading to higher levels of expression in fusion cells.

### Blimp-1 regulates a variety of genes involved in tube maturation

To further study the influence of *Blimp-1* in tracheal development, we searched *in silico* for Blimp-1 binding sites in the promoter regions of all *D. melanogaster* genes using the Matscan software (Blanco *et al.* 2006) and the reported position weight matrix corresponding to Blimp-1 (Kuo and Calame 2004). We found 3949 genes with at least one putative binding site (binding score larger than 75% of maximum value) within 2000 bases of their annotated TSSs. We prioritized candidate positions on the basis of binding score and evolutionary conservation from the UCSC *Drosophila-related* species track (Fujita *et al.* 2011) [Supplementary material T1]. In particular, we found that Blimp-1 can potentially regulate its own transcription and also the transcription of a variety of genes involved in tracheal tube development. We asked whether Blimp-1 regulation of tracheal genes was enriched in relation to all genes in the fly genome. The transcription factor Trachealess (Trh) is among the first genes to be expressed in the cells that will form the trachea. In the absence of Trh, tracheal cells fail to invaginate to form tubes and remain at the embryo surface (Isaac and Andrew 1996; Wilk *et al.* 1996). It has been shown that expression of nearly every tracheal gene requires *trh* (Chung *et al.* 2011). Thus, we wondered how many genes regulated by Blimp-1 were also downstream of Trh and, therefore, bonafide tracheal genes. To test this we combined published tracheal gene sets (Chung *et al.* 2011; Hammonds *et al.* 2013) and checked for enrichment among the genes with predicted Blimp-1 binding sites. We found that for a conservation threshold of 2 and Matscan scores larger than 75% tracheal genes are significantly enriched (fisher-test OR=1.60, p-value<0.001). In order to show robustness against the choice of Matscan score threshold we repeated the Fisher test for values ranging from 75% to 90%. We found that the odds ratio was kept relatively constant until 84%, dropping to non-significant values for the remaining thresholds (Fig. 3 B, C and Supplementary material T3).

**Figure 3.**
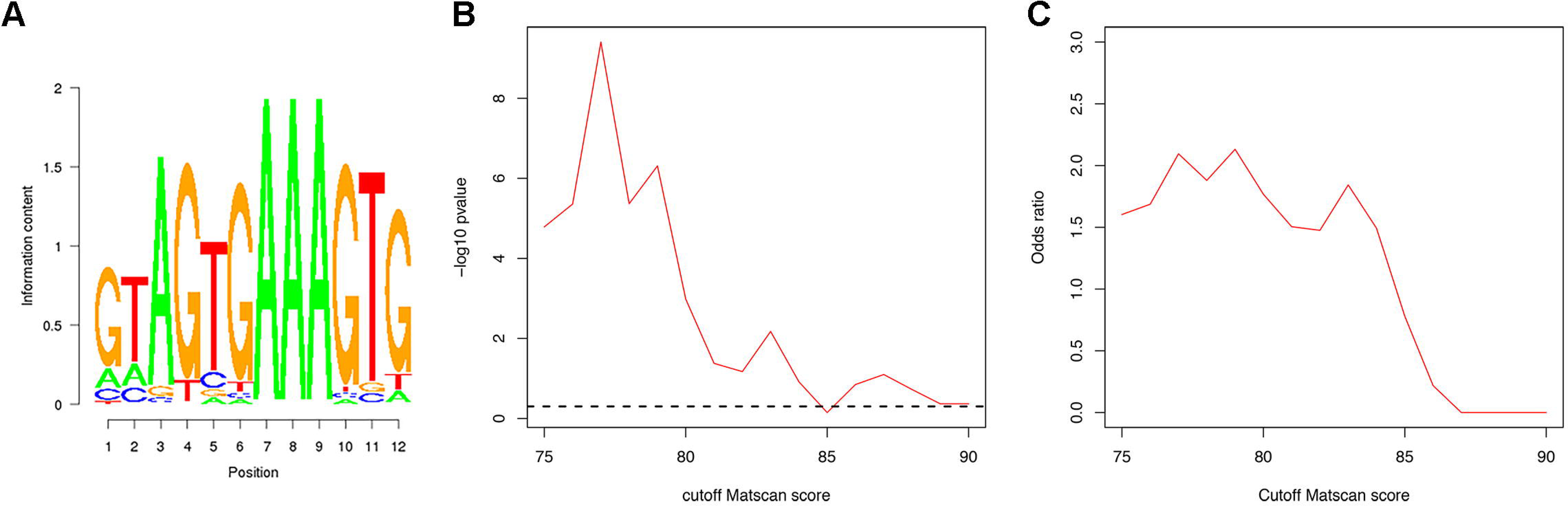
Blimp-1 position weight matrix. (A) Position weight matrix used in this study, which was taken from the reported binding sequences in (Kuo and Calame 2004). (B, C) Enrichment analysis of known tracheal genes among genes predicted to be regulated by Blimp-1. P-values and odd-ratios are shown for Fisher tests with varying Matscan scores for the definition of predicted binding sites. For values below 85 odd-ratios are relatively constant and larger than 1.5 (associated p-values < 0.05, dashed line) showing robustness against variations of the Matscan parameter.

We then selected a shorter list of genes reported to be involved in tube maturation stages (Table 1).

**Table 1.** Possible downstream targets of Blimp-1 known to be involved in tracheal tube maturation. Matscan score within the upper 75 percentile (and conservation score larger than 2).

| **Gene** | **Function** | **source** |
| --- | --- | --- |
| *atypical Protein Kinase C (aPKC)* | Member of the Par complex | (HOSONO *et al.* 2015) |
| *bitesize (btsz)* | Synaptotagmin-like protein | (JAYANANDANAN *et al.* 2014) |
| *Blimp-1* | Transcription factor | (CHAVOSHI *et al.* 2010) |
| *Btk family kinase at 29A (Btk29A)* | Non-receptor tyrosine kinase | (TSIKALA *et al.* 2014) |
| *expansion (exp)* | Required for chitin deposition | (MOUSSIAN *et al.* 2015) |
| *gartenzwerg (garz)* | Secretion | (CHUNG *et al.* 2011) |
| *grainy-head (grh)* | Transcription factor | (HEMPHÄLÄ *et al.* 2003) |
| *knickhopf (knk)* | Chitin organization | (CHUNG *et al.* 2011) |
| *obstructor-A (obst-A)* | Chitin-binding protein | (CHUNG *et al.* 2011) |
| *pebbled (peb)* | Transcription factor | (WILK *et al.* 2000) |
| *rebuf (reb)* | Required for chitin deposition | (MOUSSIAN *et al.* 2015) |
| *shade (shd)* | Ecdysone pathway enzyme | (CHAVOSHI *et al.* 2010) |
| *sinuous (sinu)* | Septate junction component | (WU *et al.* 2004) |
| *tramtrack (ttk)* | Transcription factor | (ARAÚJO *et al.* 2007) |

Clearly, this is not an exhaustive list as it only includes the genes detected using these restrictive parameters. Thus, for example, this analysis did not allow us to detect any Blimp-1 binding sites in the *tal/pri* region. However, due to the changes in *tal/pri* expression in *Blimp-1* mutants, we further studied the *tal/pri* region under less restrictive conditions. We detected four binding sites with Matscan scores above the 72 percentile and conservation scores of 1.

### Blimp-1 regulates the levels of Exp, Reb, aPKC, Knk and Btk29A

In order to further analyze the relationship between *Blimp-1* and tracheal maturation, we compared the levels of five key proteins, Exp, Reb, aPKC, Knk and Btk29A, in tube maturation in the heterozygous and homozygous *Blimp-1* mutant embryos. Exp and Reb are atypical Smad-like proteins that regulate tube size in the tracheal system by promoting chitin deposition (Moussian *et al.* 2015). aPKC is a serine/threonine protein kinase required for apico-basal cell polarity and a member of the Par complex. It has been shown to be involved in the orientation of actin rings and taenidial ridges in larval stages of tube maturation (Hosono *et al.* 2015). Knk is a GPI anchored protein needed for chitin organization and the regulation of tracheal tube diameter (Moussian *et al.* 2006). Btk29A (also known as Tec29A) is the only member of the Tec family of kinases in *Drosophila*, and it is expressed in many developmental stages of the fly. In the tracheal system, Btk29A is involved in spiracular chamber invagination, as well as in tracheal cuticle patterning (Matusek *et al.* 2006; Tsikala *et al.* 2014). Mutants and overexpression conditions for each of these genes showed phenotypes that can be correlated to the *Blimp-1* tracheal maturation phenotypes. We therefore hypothesized that Blimp-1 acts as a transcriptional repressor of these genes during tube maturation (Matusek *et al.* 2006; Moussian *et al.* 2006; Hosono *et al.* 2015; Moussian *et al.* 2015).

At early tube maturation stages, *Blimp-1* mutants displayed higher levels of Exp, aPKC, Knk and Btk29A than same-stage *wt* embryos (Fig. 4), thereby suggesting that Blimp-1 represses the expression of these proteins, as hypothesized. Of the four proteins studied, Btk29A levels showed a more pronounced difference between heterozygous and homozygous *Blimp-1* mutant embryos. Btk29A levels at stage 14 were hardly detectable in the tracheal system of *wt* embryos (Fig 4 G’)(Tsikala *et al.* 2014), whereas in mutant tracheal cells they were detected at much higher levels (Fig 4 H’). Taken together with our previous *in silico* approach, our findings suggest that Blimp-1 directly regulates the levels of these five proteins in tracheal cells during tube maturation, working as a repressor during early tracheal developmental stages.

**Figure 4.**
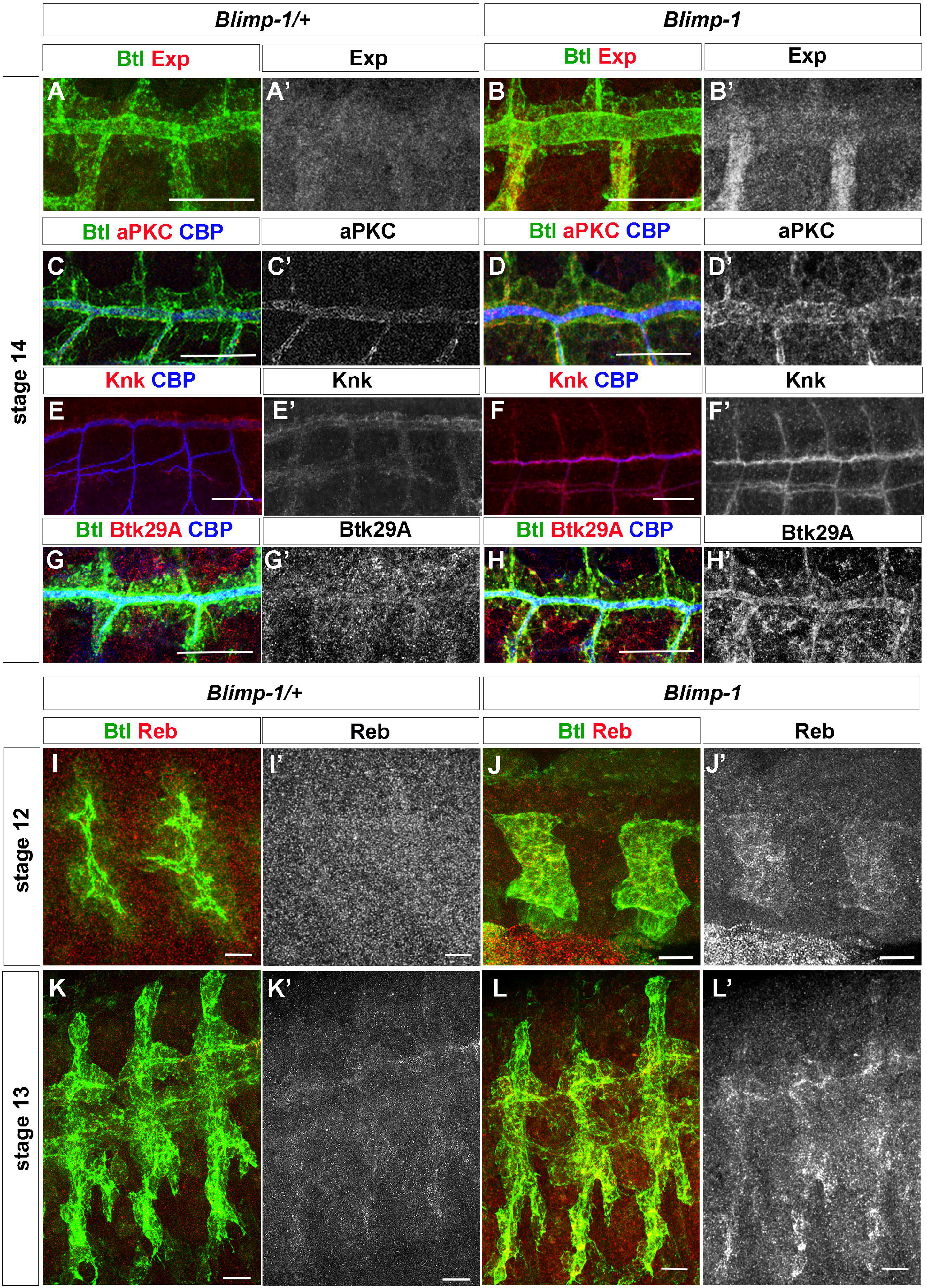
Blimp-1 regulates the expression of genes involved in tube maturation. (A,B) Exp, (C,D) aPKC, (E,F) Knk and (G,H) Btk29A levels are higher in *Blimp-1* mutants than in the *wild-type* at stage 14. Scale bars are 20 μm. (I-L) Reb expression starts earlier in *Blimp-1* mutant embryos; it is already detected in stage 12 mutants whereas it starts at stage 13 in heterozygous *Blimp-1* embryos (I,J); heterozygous stage 12 embryos in panel I are older than mutant embryos in panel J reinforcing the earlier expression of Reb in *Blimp-1* embryos. (K,L) Reb expression is higher in stage 13 *Blimp-1* mutant embryos. Scale bars are 10 μm.

During lumen maturation, chitin deposition requires the activity of Krotzkopf verkehrt (Kkv) together with Exp and Reb (Moussian *et al.* 2015). However, we could not find any evidence that Blimp-1 regulates *kkv* expression. Due to the earlier chitin deposition observed in *Blimp-1* mutant embryos (Fig.2), we wondered if Reb expression was also detected earlier, consequently leading to earlier chitin deposition. Indeed, we observed that in Blimp-1 homozygous embryos, Reb expression was already detected at stage 12, in contrast to wildtype and Blimp-1 heterozygous embryos where it starts being detected at stage 13 (Fig. 4 I-L) (Moussian *et al.* 2015). Thus, the earlier chitin deposition detected in Blimp-1 embryos may be triggered by this earlier expression of Reb together with the higher levels of Exp in tracheal cells.

### Btk29A works downstream of *Blimp-1* to regulate luminal aECM organization

What is the role of Btk29A in tube maturation downstream of Blimp-1? At late embryonic stages, a strong *Btk29A* mutant allele displays disorganized apical F-actin bundles and taenidial ridges (Matusek *et al.* 2006; Ozturk-Qolak *et al.* 2016), with heterogeneous patterns of F-actin bundling in the same trachea, showing stretches of perpendicular bundles followed by stretches of parallel bundles (Ozturk-Qolak *et al.* 2016). In addition, overexpression of full-length *Btk29A* in all tracheal cells gave rise to an expansion phenotype similar to *Blimp-1* mutants at stage 16 (Fig. 5 C, compare to 1 H). Thus, and due to our hypothesis of *Blimp-1* being a repressor *of Btk29A* expression, we examined whether derepression of Btk29A might partially account for the *Blimp-1* mutant phenotype. To do so, we used *Btk29A^K00206^*, a hypomorphic allele in which low mRNA levels are still detected in the embryonic tracheal system (Tsikala *et al.* 2014). In this hypomorphic allele, there were no apparent differences in chitin deposition between heterozygous and homozygous *Btk29A* mutant embryos at stage 16 (Fig. 5 A-B). At late stage 17, *Btk29A ^K00206^* homozygous embryos showed mild DT phenotypes compared to heterozygous embryos, such as a very mild DT expansion phenotype (Fig. 5 D, E). However, there were no detectable taenidial ridge orientation phenotypes in stage 1*7 Btk29A^K00206^* mutants, which showed parallel taenidial ridges perpendicular to the tube length as in the wild-type (Fig. 5 E INSET?). We then combined the *Btk29A^K00206^* mutation with *Blimp-1* and observed that the incomplete removal of embryonic *Btk29A* could partially rescue the taenidial ridge orientation phenotypes in most of the embryos examined (64%, n=11). These double *Btk29A^K00206^; Blimp-1* mutants showed heterogeneous patterns of taenidial ridge organization in the same trachea, showing stretches of perpendicular bundles followed by stretches of parallel ones (45% of embryos, Fig. 5 G, H, arrow in G), as well as diagonal ridges (19% of embryos, Fig. 5 H, arrow). We also observed that *Btk29A; Blimp-1* double mutant DTs showed a milder tube expansion phenotype than *Blimp-1* mutants (Fig. 5 F-H). Taken together, these results indicate that part of the *Blimp-1* phenotype can be attributed to excess *Btk29A.*

**Figure 5.**
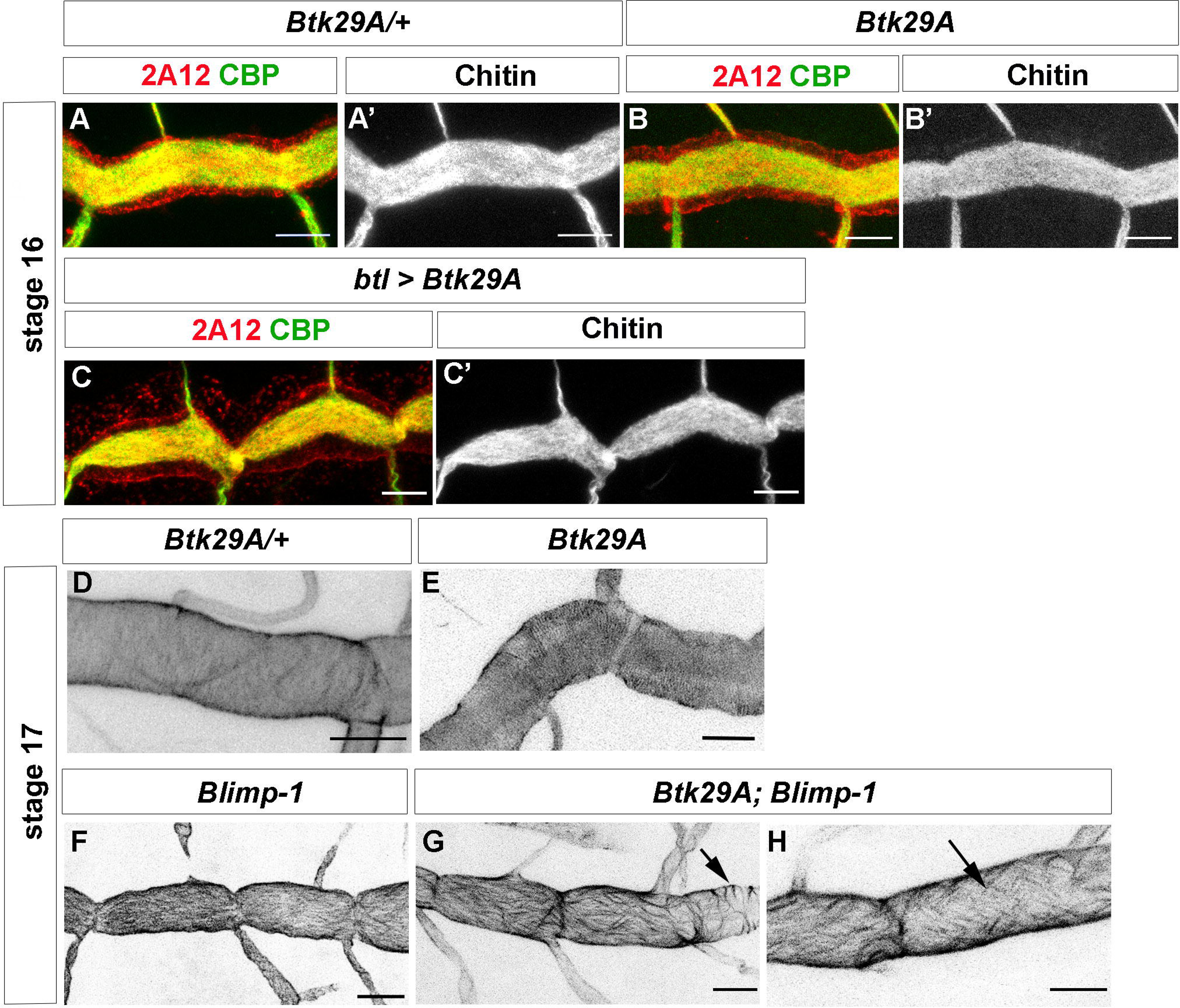
*Btk29A* partially rescues *Blimp-1* phenotypes. (A,B) *Btk29A^K00206^* embryos do not show any strong tube maturation phenotypes at late embryonic stages. (C,D) *Btk29A* overexpression in tracheal cells induces tracheal phenotypes similar to *Blimp-1* mutant embryos. (D-H) At the end of embryogenesis, stage 17, taenidial ridges can be detected by staining with chitin binding probes; *Btk29A ^K00206^* embryos display mild taenidial ridge phenotypes (E); however, when in a *Blimp-1* mutant background, a *Btk29A* hypomorphic mutation such as the one present in the K00206 allele, partially rescues the *Blimp- 1* expansion and taenidial ridge phenotypes (G,H). Scale bars are 5 μm in all panels.

## Discussion

Here we found that *Blimp-1* regulates multiple tracheal targets, thus acting as a key gene in tracheal development (Table 1 and Fig. 6A). *Blimp-1* is an ecdysone response gene (Beckstead *et al.* 2005; Chavoshi *et al.* 2010) (Fig. S1) and therefore a link between the hormonal signal and the timing of tracheal tube maturation in both embryos and larvae. We show that *Blimp-1* regulates the expression of many genes required for tube maturation. Interestingly, *in silico*, we detected four Blimp-1 binding sites in *Blimp-1* regulatory sequences using the parameters described, which suggest that Blimp-1 may regulate its own expression. This is in agreement with recent data showing that Blimp-1/PRDM1 is also able to regulate its own expression in mammals (Mitani *et al.* 2017). Self-regulation of expression is consistent with the feedback loops in which Blimp-1/PRDM1 participates and also with its role in regulating many developmental processes (Gong and Malek 2007; Bikoff *et al.* 2009). We also found Blimp-1 binding sites in the region of Tramtrack (Ttk), another transcription factor involved in many features of tube maturation (AraÚjo *et al.* 2007).

**Figure 6.**
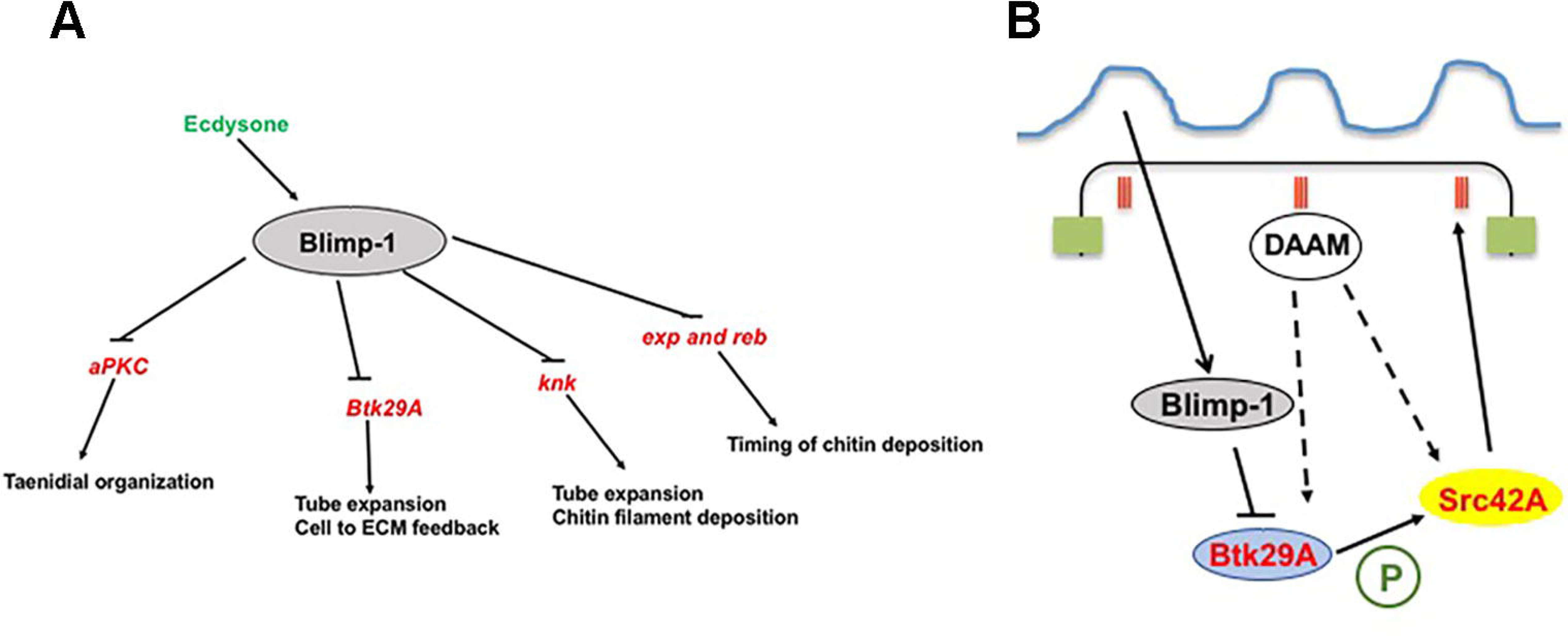
A model for the involvement of *Blimp-1* in tracheal tube maturation timing. (A) Under the influence of the insect hormone ecdysone, Blimp-1 regulates tracheal tube maturation timing by inhibiting the expression of the key genes for tube maturation, among them *aPKC, Btk29A, knk, exp and reb.* Some of these targets are known to be activated by other transcription factors like, for example Ttk in the case of Exp, Spalt (Sal) in the case of Reb and Trh in the case of Knk (Chung *et al.* 2011; Moussian *et al.* 2015). (B) Blimp-1 affects taenidial ridge formation by acting as a transcriptional repressor of Btk29A, which is an important molecule for Src42A phosphorylation. Phosphorylated Src42A and the formin DAAM regulate the actin cytoskeleton inside the cells, hence giving rise to taenidial ridges at the extracellular matrix.

Furthermore, we observed that *Blimp-1* modulates the timing of the expression of *Reb* and *Exp*, two genes involved in the genetic programme triggering timely chitin deposition (Moussian *et al.* 2015). Untimely chitin deposition was shown to disturb tube maturation, thereby demonstrating that this process has to be tightly regulated during tracheal development. Tracheal overexpression of *reb* leads to earlier chitin deposition in all branches from stage 13 and sometimes chitin appearance at stage 12 ((Moussian *et al.* 2015) and M. Llimargas personal communication). Accordingly, our results show that Reb is expressed earlier in *Blimp-1* embryos (Fig. 4 J, L). This agrees with the early chitin deposition phenotypes observed in *Blimp-1* mutants (Fig.2 B, C). Furthermore, *Blimp-1* also modulated *knk* expression during tube maturation stages. Knk is involved in directing chitin assembly in the trachea (Moussian *et al.* 2006) and correct amounts of Knk at specific times during metamorphosis are important for correct wing cuticle differentiation and function (Li *et al.* 2017). Taken together, these *in silico* and *in vivo* results indicate that Blimp-1 is a transcription factor that acts downstream of ecdysone and that it is involved in the correct timing of chitin synthesis and deposition during embryonic development.

We also found Blimp-1 binding sites in the aPKC coding region. aPKC is involved in the junction anisotropies that orient both actin rings and taenidial ridges in the lumen of tracheal tubes (Hosono *et al.* 2015). In *Blimp-1* mutants, both actin rings and taenidial ridges are either undetectable or misoriented (Ôztürk-Çolak *et al.* 2016)—observations that are consistent with changes in junction anisotropy.

We previously showed that Blimp-1 regulates chitin deposition levels and architecture and that subsequently the chitin aECM feeds back on the cellular architecture by stabilizing F-actin bundling and cell shape via the modulation of Src42A phosphorylation levels (ÖztÜrk-Çolak *et al.* 2016). However, in this report, we provided no link between the chitinous aECM and Src42A. *Btk29A* mutant larvae have an aECM phenotype, which may be the result of their actin bundle phenotype (Matusek *et al.* 2006; ÖztÜrk-Çolak *et al.* 2016). Here, we found that *Btk29A* removal can partially rescue the *Blimp-1* taenidial orientation and expansion phenotype. In view of these results, we propose that the contribution of Btk29A can be added to the feedback model for the generation of supracellular taenidia put forward in Ôzturk-Çolak et al. (Ôztürk-Çolak *et al.* 2016). We now add on our previous model by hypothesizing that Blimp-1 acts as a link between the aECM and cells by regulating the levels of Btk29A (Fig. 6 B). Btk29A and Src42A, together with the formin DAAM, have been shown to regulate the actin cytoskeleton (Matusek *et al.* 2006). In agreement with our results, we speculate that Btk29A might phosphorylate Src42A and that this phosphorylation event could be modulated by Blimp-1 and DAAM.

To conclude, our results indicate that Blimp-1 is a key player in the regulation of tracheal tube maturation and, consequently, in the feedback mechanism involved in the generation of supracellular taenidia.

## Acknowledgements

We thank M. Llimargas, E. Sanchez-Herrero, S. Roy, H. Ueda and the Bloomington Stock Center for flies and reagents. Thanks also go to the IRB Barcelona Biostatistics and Bioinformatics Unit. We acknowledge L. Bardia, A. Lladó and J. Colombelli from the IRB Barcelona Advanced Digital Microscopy Facility for help and advice with confocal microscopy and software and E. Fuentes and N. Martin for technical assistance. S.J.A. was a Ramon y Cajal Researcher (RYC-2007-00417); A.O. was the recipient of an IRB-La Caixa fellowship. This work was supported by grants from the *Generalitat de Catalunya* and the Spanish *Ministerio de Ciencia e Innovación* (BFU2009-07629).

## Supplementary material

Supplementary materials consist of the complete lists of possible genes regulated by Blimp-1 and full information on the possible binding sites as excel files and 2 Supplementary figures.

